# Competition and niche construction in a model of cancer metastasis

**DOI:** 10.1101/228585

**Authors:** Jimmy J. Qian, Erol Akçay

## Abstract

Niche construction theory states that not only does the environment act on populations to generate Darwinian selection, but organisms reciprocally modify the environment and the sources of natural selection. Cancer cells participate in niche construction as they alter their microenvironments and create pre-metastatic niches; in fact, metastasis is a product of niche construction. Here, we present a mathematical model of niche construction and metastasis. Our model contains producers, which pay a cost to contribute to niche construction that benefits all tumor cells, and cheaters, which reap the benefits without paying the cost. We derive expressions for the conditions necessary for metastasis, showing that the establishment of a mutant lineage that promotes metastasis depends on niche construction specificity and strength of interclonal competition. We identify a tension between the arrival and invasion of metastasis-promoting mutants, where tumors composed only of cheaters remain small but are susceptible to invasion whereas larger tumors containing producers may be unable to facilitate metastasis depending on the level of niche construction specificity. Our results indicate that even if metastatic subclones arise through mutation, metastasis may be hindered by interclonal competition, providing a potential explanation for recent surprising findings that most metastases are derived from early mutants in primary tumors.

## 1 Introduction

A cancer tumor is a collection of abnormal cells whose unregulated proliferation damages surrounding host tissue, often resulting in patient death. It is also a population of genetically and phenotypically diverse cells that compete, propagate, and contribute (or not) to the cellular society. Tools from population biology are therefore increasingly used to study cancer dynamics. Cancer’s genetic instability and high mutation rate, compounded with harsh spatial constraints, a dearth of nutrients, and immune surveillance, lead to rapid selection for the survival of the fittest tumor cells. However, the evolutionary dynamics of tumors are only fully comprehensible when the ecological context – the tumor ecosystem – is considered [1, 2]. This entails applying ecological concepts such as predation, niches, and invasion (in evolutionary theory and in this paper, “invasion” refers to the establishment of a mutant genotype into an existing population, a concept distinct from cancer “invasion,” or expansion, into surrounding tissue). Accordingly, a number of ecological models have provided useful insight into cancer progression [3, 4].

A recently influential idea in ecology is that not only does the environment act on a population to generate selection pressures and Darwinian evolution, but organisms reciprocally modify the environment through a process called niche construction (also known as ecological engineering) [5, 6]. Via niche construction, organisms not only influence aspects of the ecosystem such as resource flow and trophic relationships, but they modify the actual sources of natural selection acting on themselves and their neighbors. For example, new selection pressures on beavers’ teeth, tail, and social behavior arise due to the construction of a dam [6]. The environmental modifications resulting from niche construction may be passed down to descendants through ecological inheritance, which has been recognized as a key aspect of extra-genetic inheritance [7].

Niche construction also likely plays an important role in cancer population biology [8–11]. Cancer cells greatly alter their microenvironments. For example, tumor cells release angiogenic factors such as vascular endothelial growth factor and stimulate vascularization [12–14], reduce local pH [15], release a gamut of growth factors such as insulin-like growth factor II [16], and secrete matrix metalloproteinases that degrade extracellular matrix proteins [12]. Tumors also drastically alter the local flow of nutrients and signaling factors, creating a nutrient-poor ecosystem that is passed down to descendant cells via ecological inheritance. This ecological inheritance promotes tumor cell heterogeneity and cancer growth, suggesting that cancer niche construction may be a worthwhile therapeutic target [8].

In this paper, we use niche construction theory to examine metastasis. Metastasis is not simply a result of mutation of tumor subclones into more invasive phenotypes and subsequent cell dissemination; it additionally requires the construction of a pre-metastatic niche [10, 17–22]. The concept of the pre-metastatic niche dates back to Paget’s “seed and soil” hypothesis, which states that tumors (the “seed”) are predisposed to metastasize to certain organs (the “soil”) because the metastatic site must provide a milieu conducive to the recruitment and settlement of disseminated tumor cells [23]. This receptive microenvironment, termed the pre-metastatic niche, must be established before metastasis can occur [10, 17–22]. Examples of pre-metastatic niche construction include increasing vascular permeability and clot formation, altering local resident cells such as fibroblasts, remodeling the extracellular matrix, and activating and recruiting non-resident cells such as haematopoietic progenitor cells and other bone marrow-derived cells, which further induce many subsequent changes [17]. Interestingly, evidence has shown that primary tumors actively prepare distant organs for reception of future metastatic cells by secreting various factors and extracellular vesicles that foster pre-metastatic niche construction into the bloodstream [17–22, 24–29]. Primary tumor-derived secretions that promote pre-metastatic niche construction include TGF*β* [18, 21], TNF-*α* [18], placental growth factor [18, 22], vascular endothelial growth factor [18, 21, 22], lysyl oxidase [19], microvesicles [29], exosomes [20, 26–28], and many more [17, 18]. These findings show that some primary tumor cells sacrifice metabolic resources in order to promote successful settlement by their disseminated descendants into metastatic sites, which provides no benefit to themselves. Why such behavior is so common is an interesting question especially because the ability of a tumor to metastasize cannot evolve adaptively analogous to life-history traits, since tumors are not selected to metastasize between generations and cancer lineages are in general evolutionary dead-ends [2]. Accordingly, the ability to metastasize, when it does occur, arises as a result of local ecological dynamics of a tumor. In this paper, we are interested in the fate of primary tumor mutations that promote pre-metastatic niche construction, rather than the entire metastatic cascade or settlement into the metastatic site.

Although previous work has recognized the applicability of niche construction theory to cancer [8, 9], there are only a few formal models of the phenomenon. Among these, Bergman and Gligorijevic [10] proposed a framework to integrate experimental metastasis data with niche construction theory, with the goal of providing a predictive model that can be directly parameterized. Another model by Gerlee and Anderson [30] studied the evolution of tumor carrying capacity as a function of niche construction. They assumed that niche construction increases the tumor carrying capacity, a phenomenon commonly seen in ecological settings. They noted that tumors may include both producers, which actively contribute to niche construction, and cheaters, which reap the benefits of niche construction without paying the growth rate cost of production. They showed that the specificity of the benefits from niche construction as well as spatial structure maintains selection for producers and allows for coexistence of cheaters and producers.

Another idea that motivates our model is the recent observation that metastatic cell lineages tend to diverge from the primary tumor early on [31]. In other words, metastasis involves mutations that occur early in the tumor’s lifetime. This finding contradicts the linear progression model of cancer, where metastatic tumors arise from late-stage primary tumors. This finding is somewhat paradoxical, since later-stage primary tumors are bigger and therefore harbor more mutations from which metastatic tumors might arise, and hence one might expect more metastatic tumors to be derived from late-stage tumors. As we will see below, competition between local and pre-metastatic niche constructors may provide a potential answer to this paradox.

We present a mathematical model of niche construction and metastasis in cancer. Our model contains producers of both the primary tumor (i.e., local) niche and the pre-metastatic niche, as well as cheaters. We model a tumor population with a carrying capacity that increases with local niche construction. We derive expressions for the ecological conditions necessary for metastasis, showing that they depend on niche construction specificity and the interclonal competition structure. Our results reveal a robust trade-off between the arrival of metastasis-promoting mutants and their ability to invade a tumor. Tumors composed only of cheaters remain small but are susceptible to invasion by cells that construct the pre-metastatic niche, whereas larger tumors containing producers may be unable to facilitate metastasis depending on the level of niche construction specificity. In certain competition structures, tumors containing only local producers can completely preclude metastasis unless invasion of metastasis-promoting subclones occurs early on. Our results highlight the fact that metastasis requires both the necessary genetic mutations and a suitable ecological milieu: even if metastatic subclones arise through mutation, invasion may not be possible due to competitive exclusion and a lack of niche opportunities. These findings can explain the observation that metastasis involves early mutations [31].

## 2 Methods

We consider a primary tumor with *N* cells, which can include both producers and cheaters, and a bloodstream into which tumor cells can enter via intravasation. (An extended form of the model is shown in Supplementary Information (SI) section SI-A.) Producers participate in niche construction at a cost to their growth rate, since it takes energy and metabolic resources to secrete angiogenic factors, growth factors, and matrix metalloproteinases. Cheaters do not participate in niche construction but still benefit from it, so they have a higher growth rate than producers. We assume that a cell’s type (producer or cheater) is determined genetically.

There are three subsets of producers. Local producers contribute only to niche construction in the tumor’s immediate microenvironment, benefiting primary tumor cells but not circulating or metastasized cells. The extent of local niche construction is represented by the amount of resource *R*, a general resource that for example could represent the amount of recruited vasculature. The primary tumor also includes secondary producers, which contribute to the spatially distant pre-metastatic niche by secreting chemokines, growth factors, and exosomes into the bloodstream to allow circulating tumor cells to settle down to form a secondary tumor, as mentioned in the introduction. These molecules are carried away from the primary tumor and provide no benefit to primary tumor cells, so construction of the pre-metastatic niche is not included in the variable *R*. Secondary producers pay a growth cost similar to primary producers, but they otherwise act as cheaters from the primary tumor’s point of view since they benefit from *R* without contributing to it. Additionally, there are global producers that contribute to niche construction in both the primary microenvironment and the pre-metastatic niche and pay double the growth rate cost. Because pre-metastatic niche construction is required for metastasis, as discussed above, we treat the existence of secondary or global producers as a necessary condition for metastasis, consistent with our focus on interrogating the prerequisites of metastasis within the primary tumor. SI section SI-B discusses how our model is robust to changes in the interpretation of the four cell types and shows how our model may be generalized without changing the mathematical details or results.

Cells are given a subscript (*x,y*), where *x* ∈ {0,1} describes participation in local niche construction (0 for cheaters and 1 for producers) and *y* ∈ {0 1} similarly denotes participation in pre-metastatic niche construction. The population of each cell type is *n*_*xy*_ with respective growth rates *r*_*x,y*_. Local and global producers increase *R* with rate *g* and *R* suffers independent resource depletion with rate *l*.

**Figure 1:**
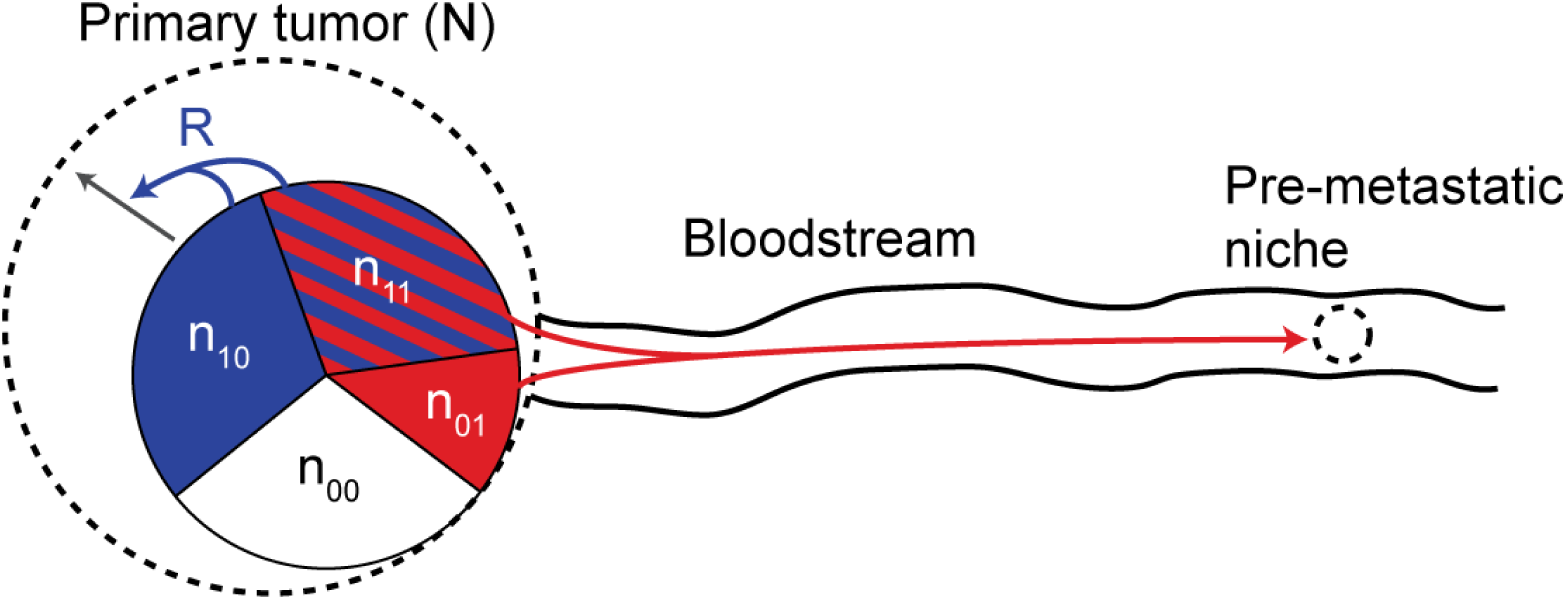
Schematic representation of the model, which considers a primary tumor with four cell types and a distant pre-metastatic niche. Cheaters are white, local producers are blue, secondary producers are red, and global producers are both red and blue. Niche construction occurs in the primary microenvironment through production of resource *R*, which benefits the tumor by increasing carrying capacity, represented a as dotted line. Construction of the pre-metastatic niche by primary tumor cells is represented by the red arrow.

Primary tumor cells enter the bloodstream as a result of intravasation. Local crowding has been suggested to cause a reduction in tumor cell fitness and lead to increased mutation rate and ecological dispersal [8]. Other studies have provided evidence that haematogenous tumor cell dissemination can begin early during primary tumor development and progression [21,32,33]. To account for these results and the ecological dispersal hypothesis, we introduce a function *m*(*N, R*) representing the rate at which primary tumor cells exit the local niche and enter the bloodstream. We assume this function has the form 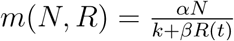 where *α* is a constant and the denominator is the carrying capacity (discussed below). Cells tend to migrate more when they receive less of the share of resources in the microenvironment. It is important to note that the precise form of this dispersal function is not crucial to our results, because parameter estimation (see SI section SI-E) suggests *²* is several orders of magnitude smaller than any other parameter, a fact we use in simplifying our results as described later.

### 2.1 Carrying capacity

We assume carrying capacity increases linearly with niche construction. Primary and secondary tumors possess intrinsic carrying capacity *k*, which can represent the number of cells that can survive without significant self-induced angiogenesis or release of growth factors. In the primary tumor, the carrying capacities of cheaters and secondary producers are both *k* + *β*_0_*R*(*t*) while those of primary and global producers are *k* + *β*_1_*R*(*t*).*β*_0_ and *β*_1_ are constants describing the benefit that either cheaters or producers receive from niche construction. If *β*_0_ ≠ *β*_1_, then either cheaters or producers use the resource more efficiently. This is analogous to the specificity of niche construction in Gerlee and Anderson’s model [30]. If 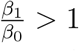, modifications of the niche are specific to the genotype that generates it and cheaters are less able to free-ride. Strong specificity refers to 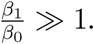

### 2.2 Competition

We assume cells grow according to Lotka-Volterra competition equations, shown in Table 1 with the parameters summarized in Table 2. We consider multiple competition structures with varying competition strength among the four cell types,summarized in Figure 2. In each, the strength of inter-type competition between the four cell types (symmetric in competition structures I and II) is denoted by Greek letters whose values are positive and less than or equal to 1. The magnitude of intraclonal competition is 1, such that interclonal competition strength is weaker than or equal to intraclonal competition. Biologically, stronger intra-type competition can stem from spatial considerations since cellular neighbors tend to be of the same cell type. Alternatively, stronger intra-type competition can arise because different cell types utilize other resources (that we do not explicitly model) differentially. For example, cheaters focus on cell division and require a significant commitment to nucleotide biosynthesis and genome duplication. Producers, on the other hand, focus on protein production.

**Figure 2:**
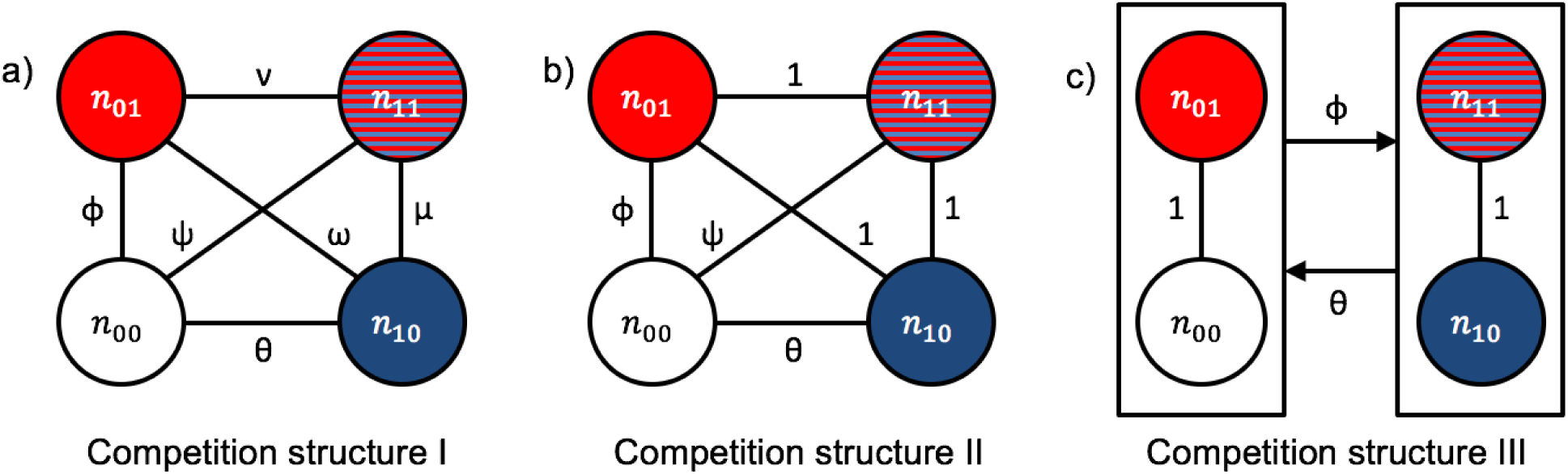
Schematic representation of the different competition structures we consider. The strength of competition between each cell type is shown along connections in the lattice. Intraclonal competition is 1 for all cell types. *ϕθ*, and *Ψ* are positive and less than 1. *v, μ*, and *ω*, are positive and less than or equal to 1. In competition structure III, the two distinct niches are represented by boxes. Cells that cheat in the primary tumor experience competition of magnitude *θ* due to, and compete with magnitude 0 with, cells that produce the primary resource.

Competition structure I is the most general symmetric case. In competition structure II, interclonal competition between producers is as strong as intraclonal competition, while cheaters compete less with all three producer types. This scenario may arise if primary and secondary niche construction require similar metabolic resources so all producers occupy the same niche, whereas cheaters focus on their own division instead of ecological engineering. Competition structure III assumes the two niches are producing and cheating in the primary tumor regardless of propensity for secondary resource production. From the primary tumor’s standpoint, cheaters and secondary producers may occupy the same niche since neither cell type participates in local niche construction, while local and global producers both do and thus occupy a distinct niche. Intra-niche competition is as strong as intraclonal competition, while inter-niche competition is weaker.

### 2.3 Separation of time-scales

Simulations of the model (shown in SI section SI-D) show that, for reasonable parameters (inferred from the literature in SI-E), cell populations equilibrate more quickly than the resource dynamics. The latter keep growing without reaching an equilibrium at time-scales relevant to tumor growth (i.e. the lifespan of a human). This is biologically intuitive since niche construction processes such as microenvironment vascularization are generally slower than cell division. This allows us to make a separation of time-scales argument. In particular, we consider the cell dynamics (equations 11-1.4) to be fast and the resource dynamics (equation 1.5) to be slow We first analyze the fast-changing variables while treating the slow-changing variable as constant. In other words, we find the equilibria of equations 1.1-1.4 while holding *R* constant (we refer to these equilibria of the fast dynamics, which are functions of *R*, as “quasi-equilibria”). Then, we analyze the dynamics of the slow variable *R* while assuming the fast variables are at a quasi-equilibrium.

**Table 1:** Governing equations of the model under competition structure I, and the corresponding variables whose rates of change they describe. Time dependence of *n* and *R* have been suppressed for notational simplicity. Dependence of *m* on *N* and *R* has also been suppressed. The equations for competition structures II and III are shown in SI section SI-C.

|  |  |  |
| --- | --- | --- |
| Primary cheaters | $\frac{dn_{00}}{dt} = r_{00}n_{00}\left(1 - \frac{n_{00} + \phi n_{01} + \theta n_{10} + \psi n_{11}}{k + \beta_0 R}\right) - mn_{00}$ | (1.1) |
| Secondary producers | $\frac{dn_{01}}{dt} = r_{01}n_{01}\left(1 - \frac{\phi n_{00} + n_{01} + \omega n_{10} + \nu n_{11}}{k + \beta_0 R}\right) - mn_{01}$ | (1.2) |
| Primary producers | $\frac{dn_{10}}{dt} = r_{10}n_{10}\left(1 - \frac{\theta n_{00} + \omega n_{01} + n_{10} + \mu n_{11}}{k + \beta_1 R}\right) - mn_{10}$ | (1.3) |
| Global producers | $\frac{dn_{11}}{dt} = r_{11}n_{11}\left(1 - \frac{\psi n_{00} + \nu n_{01} + \mu n_{10} + n_{11}}{k + \beta_1 R}\right) - mn_{11}$ | (1.4) |
| Resource | $\frac{dR}{dt} = g(n_{10} + n_{11}) - lR$ | (1.5) |

**Table 2:** A summary of the model parameters, some of which are estimated as described in SI section SI-E.

| Parameter/variable | Description |
| --- | --- |
| $n_{xy}, r_{xy}$ | number and growth rate of $xy$-type cells |
| $\beta_0$ | benefit from niche construction for cheaters and 2° producers |
| $\beta_1$ | benefit from niche construction for local and global producers |
| $k$ | intrinsic carrying capacity |
| $\alpha$ | intravasation rate |
| $\theta, \phi, \omega, \psi, \mu, \nu$ | interclonal competition terms (see Figure 2) |
| $g$ | resource production rate |
| $l$ | independent resource depletion rate |

## 3 Results

For each competition structure, we examine a primary tumor that initially consists of only cheaters and local producers. The conditions for metastasis are equivalent to the invasion conditions of secondary or global producers into this tumor, since pre-metastatic niche construction is required for circulating tumor cells to settle into a secondary site. For invasion of secondary producers, 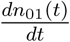 must be positive if a small but nonzero number of secondary producers cells are suddenly added to the population (e.g. through mutation). For invasion of global producers, 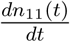 must be positive if a small but nonzero number of global producers are suddenly added to the population. There are three possible non-trivial quasi-equilibria of a local tumor: cheaters only, local producers only, and coexistence. We determine the stability of each quasi-equilibrium and evaluate the invasion conditions for secondary and global producers. These results for each competition structure are outlined in Table 3 and considered in detail below.

**Table 3:** A comparison of the invasion conditions at and stability conditions of each quasi-equilibrium for each competition structure. The conditions for stability of coexistence are omitted because they are mathematically intractable, though numerical analysis showed stability can be easily achieved for various parameter combinations. It is assumed that at the producer-only and coexistence quasi-equilibria, *R ≫ k* while at the cheater-only quasi-equilibrium, *k ≫ R*.

| Competition structure | I | II | III |
| --- | --- | --- | --- |
| Producer-only stability | $\theta > \frac{\beta_0}{\beta_1}$ | $\theta > \frac{\beta_0}{\beta_1}$ | $\theta > \frac{\beta_0}{\beta_1}$ |
| Invasion of 2° producers | $\omega < \frac{\beta_0}{\beta_1}$ | $\beta_0 > \beta_1$ | $\theta < \frac{\beta_0}{\beta_1}$ |
| Invasion of global producers | $\mu < 1$ | false | false |
| Cheater-only stability | false | false | false |
| Invasion of 2° producers | true ($\phi < 1$) | true ($\phi < 1$) | $r_{01} > r_{10}$ |
| Invasion of global producers | true ($\psi < 1$) | true ($\psi < 1$) | true ($\phi < 1$) |
| Coexistence stability | [messy] | [messy] | [messy] |
| Invasion of 2° producers | $\beta_1(\phi\theta - \omega) + \beta_0(\omega\theta - \phi - \theta^2 + 1) > 0$ | $\frac{\phi + \theta(\theta - 1) - 1}{\phi\theta - 1} > \frac{\beta_1}{\beta_0}$ | false |
| Invasion of global producers | $\beta_0(\mu\theta - \psi) + \beta_1(\psi\theta - \mu - \theta^2 + 1) > 0$ | $(\psi - \theta)(\theta\beta_1 - \beta_0) > 0$ | $(\phi - \theta)(\beta_1\theta - \beta_0) > 0$ |

Tumors containing producers have a large amount of resource, i.e. *R ≫ k*, since the producer-only and coexistence quasi-equilibria result in rapid resource accumulation. On the other hand, tumors starting with cheaters only have low *R*, i.e. *k ≫ R* since there is no niche construction. Additionally, *α* ≈ 0 in any sum since *α* is several orders of magnitude smaller than any other parameter (see SI section SI-E). We use these facts in simplifying the derivation of stability and invasion conditions. The trajectories the tumor can undergo depend on niche construction specificity and inter-type competition structure. We consider each possibility in detail below.

### 3.1 Competition structure I

The invasion conditions for secondary producer and global producers are, respectively,

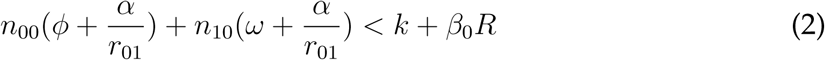

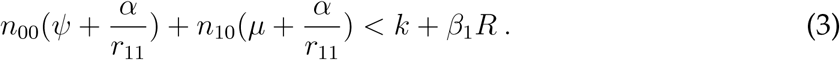

For large *R* and small *α*, the producer-only quasi-equilibrium is stable when

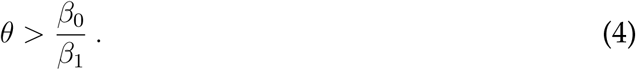

This inequality means that the higher the specificity of niche construction (measured by 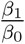) the less likely cheaters are able to invade the population. Secondary producers can invade the local producer-only tumor if

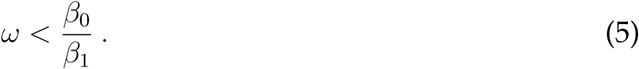

This condition similarly means that the higher the niche construction specificity, the less likely the invasion of secondary producers. If *ω* < *θ*, there is a window of specificity where the producer-only tumor is resistant to invasion by cheaters but susceptible to invasion by secondary producers. On the other hand, global producers can invade the producer-only tumor if

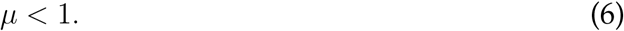

Thus, the stability and resistance to invasion of a tumor containing only producers depends on the strength of interclonal competition and may depend additionally on niche construction specificity.

Secondary producers can invade the coexistence quasi-equilibrium if

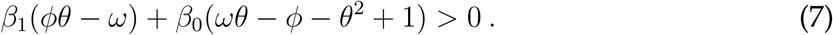

For high niche specificity, this is satisfied when *ϕθ* > *ω* which is unlikely given that competition between cheaters and any producer type is less than competition among producers. Global producers can invade a tumor at coexistence if

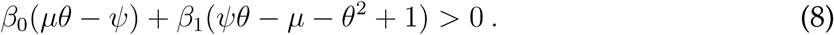

Importantly, whether a metastasis-promoting subclone can invade a tumor containing both cheaters and local producers depends on both niche construction specificity and competition strength. Finally, it is easy to see from equations 2 and 3 that a cheater-only tumor (with *R* = 0 and *α* ≪ 1) can always be invaded by any producer cell type regardless of specificity, as long as *ϕ* and *ψ* < 1.

These results point to an interesting trade-off: cheater-only tumors ofer no competitive obstacle to metastasis. However, they remain small due to the lack of niche construction, which constrains the number of mutations they might experience that can lead to secondary or global producer clones. In contrast, if local producers invade first the tumor grows bigger, increasing the arrival rate of mutations, yet simultaneously the invasion conditions for a secondary or global producer become more stringent so that the pre-metastatic niche may be precluded by competition. As we discuss below, this tension is even more apparent in other competition structures. Figure 4 schematically illustrates this trade-off for all competition structures.

**Figure 4:**
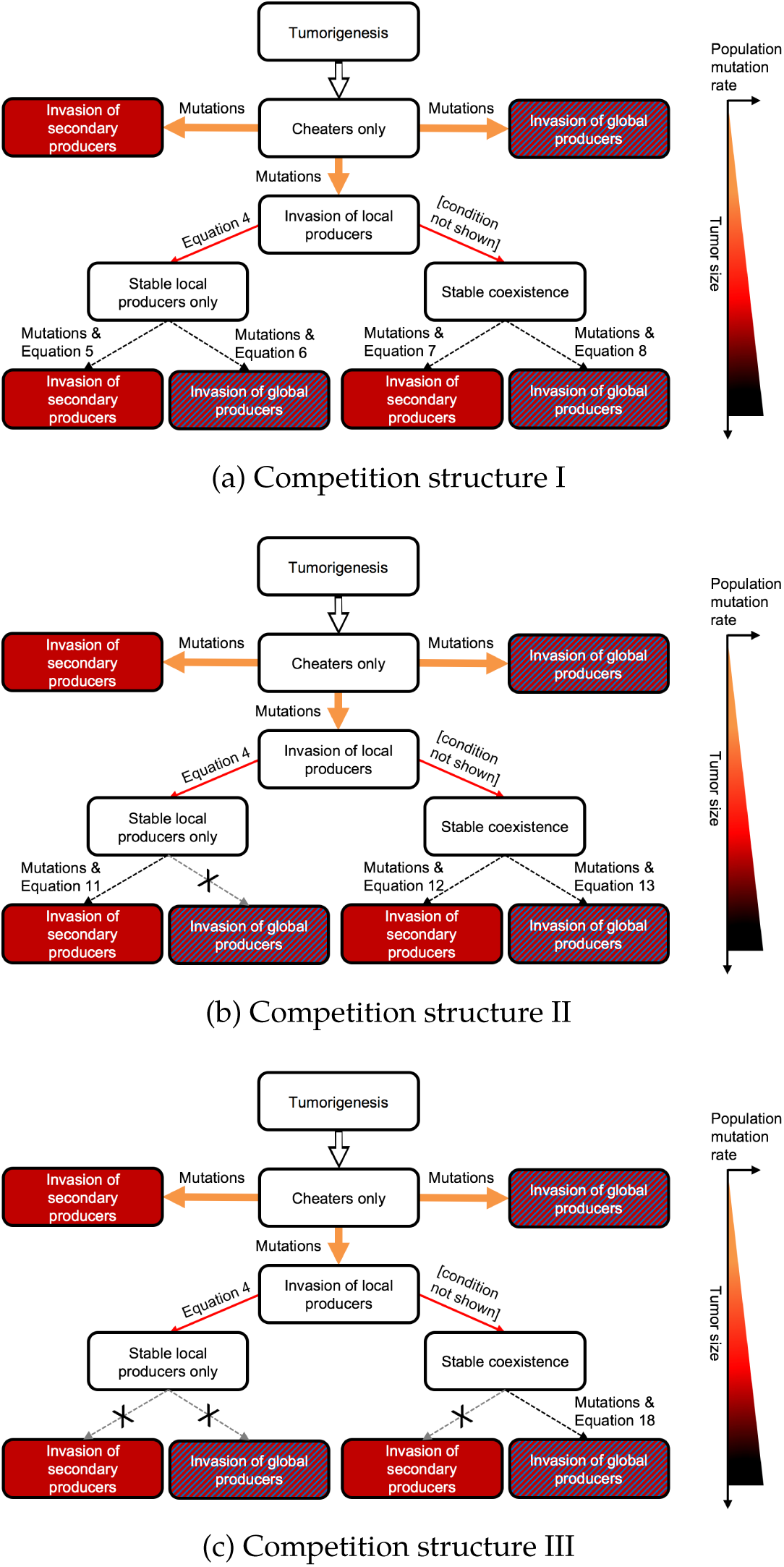
Schematic of possible tumor trajectories with their corresponding conditions. The thicker the arrow, the easier the ecological conditions are met. Arrow colors correspond to the mutation rate according to the mutation gradient on the right. Crossed out arrows indicate resistance to invasion. Tumor size and population mutation rate increase going down the flowchart, as indicated by the graph on the right.

### 3.2 Competition structure II

The invasion conditions for secondary and global producers are, respectively,

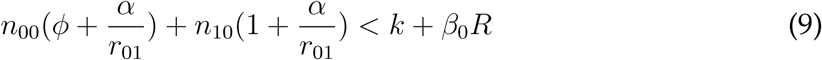

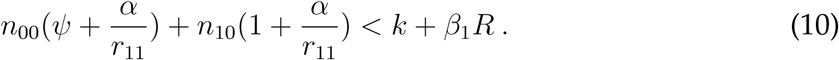

Producer-only tumors can be invaded by secondary producers when

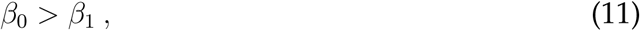

i.e. when there is no niche specificity and cells that do not produce the resource must benefit from it more than cells that do. Global producers cannot invade the producer-only tumor under this competition structure.

At the coexistence quasi-equilibrium, invasion of secondary producers can occur if

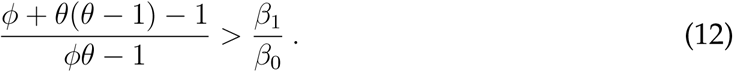

This condition is less likely to be true with increasing specificity. Global producers can invade when

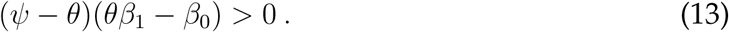

Both invasion conditions for tumors with coexistence depend on the strength of competition and niche specificity. The cheater-only tumor, on the other hand, is always vulnerable to invasion by any producer cell types, just like for competition structure I. The trade-off between mutant arrival and invasion is reproduced in this competition structure and is even more apparent since global producers cannot invade producer-only tumors. Once again, stability and invasion of tumors containing producers depend on competition strength and specificity while cheaters are generally susceptible regardless of specificity.

### 3.3 Competition structure III

The invasion conditions for secondary producers and global producers are, respectively,

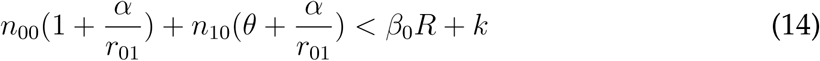

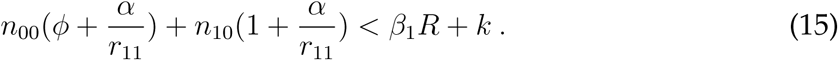

The condition for stability of the local producer-only quasi-equilibrium is equation 4, just like the other two competition structures. Global producers cannot invade producer-only tumors, while invasion of secondary producers is possible when

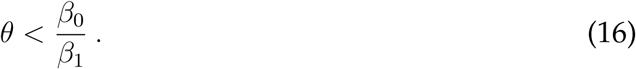

As niche construction specificity increases, this condition is less likely to be true. This invasion condition is mutually exclusive with the stability of the quasi-equilibrium. If 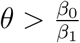 then the tumor remains at the stable producer-only quasi-equilibrium and is resistant to invasion by cheaters, global producers, and secondary producers. If 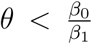 the quasi-equilibrium is unstable and susceptible to invasion by cheaters or secondary producers. The larger the competition that secondary producers would experience from local producers, the more efficiently they must be able to use the resource in order to invade.

At the coexistence quasi-equilibrium, the invasion condition for secondary producers is

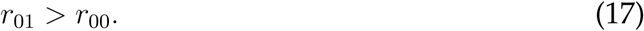

This condition is never fulfilled since secondary producers pay a growth rate cost relative to cheaters. The condition for invasion of global producers is

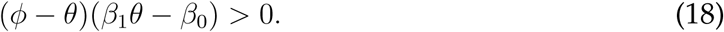

The coexistence quasi-equilibrium allows for invasion of cells that contribute to the pre-metastatic niche only if they also contribute to local niche construction and only under certain levels of interclonal competition and specificity. Even if the necessary mutations for genesis of secondary producers occur, ecological conditions prevent the invasion of the lineage. Coexistence of cheaters and local producers can obstruct successful metastasis through a failure of settlement into the pre-metastatic niche rather than a failure of intravasation.

On the other hand, the cheater-only quasi-equilibrium is unstable and always vulnerable to invasion by global producers. Secondary producers can invade if

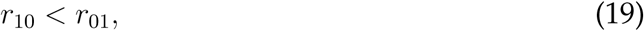

i.e. if primary producers grow more slowly than secondary producers, which may be satisfied since the growth rate cost of local niche construction can easily be higher than that of preparing the pre-metastatic niche, again highlighting the susceptibility of cheater-only tumors to invasion by all producers.

In short, for all competition structures we consider, tumors with cheaters only are easily invaded while tumors containing producers are more difficult to invade, with restrictions on competition strength and niche construction specificity. To confirm this tension between invasion and mutation, we simulated the tumor and resource dynamics starting with cheaters only. The mutation rate in cancer is estimated to be 2 × 10^−7^ per cell division per gene [34] and the cell cycle length is approximately one day for at least some cancers [35]. We thus use a daily mutation rate of 2 × 10^−7^ and assume 1 out of 1000 mutations creates (11) cells from (00) cells or (10) cells from (01) cells. We assume 1 out of 500 mutations creates (01) or (10) cells from (00) cells, or (11) or (00) cells from (10) cells, since these cellular transformations do not require as drastic a phenotypic alteration. These mutation probabilities are somewhat arbitrary but the trade-off is robust to the choice of specific mutation probabilities. We choose these specific probabilities only to illustrate this trade-off in a convenient manner.

Figure 3a shows a common tumor trajectory with clinically realistic tumor size. The tumor starts with cheaters and does not increase in size initially after arriving at the carrying capacity without producers. Once producers arise by mutation and successfully invade, cheaters go extinct. The tumor increases in size as resource production commences. The increasing size leads to numerous mutations, but these mutations do not lead to successful invasion since *R* has accumulated to a high level and we showed above that a stable producer-only quasi-equilibrium with high *R* is resistant to invasion. Figure 3b shows a clear trade-off between mutation rate and invasion. In tumors where producers arise from mutation and invade, size increases with time. The number of mutations increases drastically with tumor size, but these mutations all result in failed invasion. In tumors that remain cheater-only, successful invasion of secondary or global producers is possible, as shown by red triangles. There is a much smaller number of mutations for cheater-only tumors due to their small size, but once a mutation does arise, invasion is much more probable than in larger producer-only tumors.

**Figure 3:**
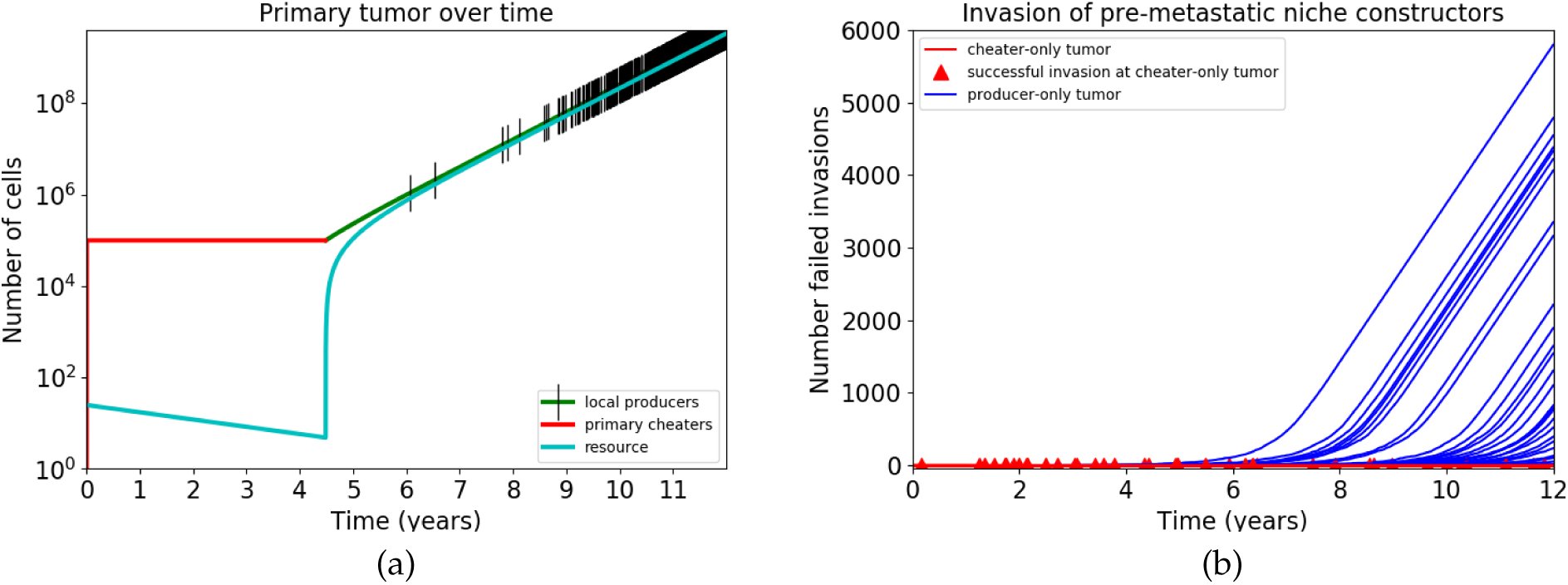
Simulation of tumors starting with cheaters. Parameters used are *τ*_00_ = 0.07, *r*_10_ = 0.05, *r*_01_ = 0.045, *r*_11_ = 0.02, *k* = 10^5^, *β*_0_ = 1, *β*_1_ = 1.2, *θ* = *ϕ* = 0.9, *g* = 0.004, *l* = 0.001, *α* = 10^−6^ some of which are estimated in SI section SI-E. Mutation rates are mentioned in the text. If successful invasion of producers occurs, cheaters become extinct rather than arrive at coexistence for these parameters. (a) Simulation of a single tumor starting with cheaters only and a small amount of resource. Black tick marks represent mutations leading to arrival of secondary or global producers, though none of them lead to successful invasion. (b) Simulation of 200 tumors starting with cheaters only. Each red triangle indicates a successful invasion of a cheater-only tumor by secondary or global producers. Each blue curve represents a tumor that has been invaded by local producers; none of these producer-only tumors experienced successful invasion by secondary or global producers despite the arrival of numerous mutants, plotted on the y-axis.

### 3.4 Tumor trajectories

Figure 4 schematically summarizes the trajectory tumors can undergo starting from cheaters only, in light of the results presented above. After tumorigenesis, cheaters proliferate and approach the intrinsic carrying capacity. The small initial tumor is always unstable and can be invaded by any producer cell type regardless of specificity. It can promote metastasis as long as the necessary mutations occur to generate secondary or global producers. However, to continue expanding the tumor population, niche construction is necessary. Mutations can lead to the appearance of producers from the cheater-only tumor, which saves the population from stagnation. Subsequent tumors reaching either coexistence or extinction of cheaters can, however, be resistant to invasion by metastasis-promoting lineages, depending on competition strength and niche construction speci-fcity. Furthermore, under competition structure III, any tumor at the stable local producer-only quasi-equilibrium is resistant for all levels of specificity and any tumor containing coexistence is resistant to invasion by secondary producers.

## 4 Discussion

We presented a simple model of niche construction in cancer, where local niche construction benefits all primary tumor cells by increasing the carrying capacity, and secondary niche construction (construction of the pre-metastatic niche) is needed for successful metastasis. Primary tumor cells can contribute to niche construction in one or both of the sites at a cost to their growth rate. Cheaters can reap the benefits of niche construction without paying the cost. Although no definitive information exists on the relative strengths of interclonal competition and density dependence, we have analyzed three plausible competition structures of varying generality.

The primary tumor, without any distant or global producers, can arrive at one of three nontrivial quasi-equilibria: extinction of cheaters, extinction of local producers, or coexistence of local producers and cheaters. The cheater-only quasi-equilibrium is vulnerable to invasion by any producer cell type, independent of niche construction specificity, as long as interclonal competition is weaker than intraclonal. On the other hand, quasi-equilibria containing producers have different requirements for stability and varying levels of susceptibility to the invasion of secondary or global producers, dependent on the strength of interclonal competition and niche construction specificity. The invasion of primary tumor cells that contribute to the pre-metastatic niche is a necessary condition for metastasis and settlement of the secondary tumor site [17–22]. Importantly, susceptibility or resistance to invasion are not intrinsic to a tumor, but are crafted through an ecological pathological relationship between the tumor and its microenvironment. Metastasis requires the necessary mutations for the genesis of certain subclones and also an ecological milieu that facilitates invasion of these subclones. Even if the appropriate mutations occur, the cells could fail to invade and instead die off if the tumor is resistant to invasion. We have shown that such resistance is more likely to occur in tumors containing producers, which are larger and accumulate more mutations. Small, cheater-only tumors experience fewer mutations yet are more able to facilitate the successful proliferation of metastasis-promoting lineages. Although we adopt a deterministic invasion perspective (i.e., mutant lineages either increase or not depending on the invasion condition), our argument also applies to the stochastic persistence of a small mutant lineage, since all things being equal, such persistence is less likely when invasion conditions are not satisfied.

Under all three competition structures, tumors containing only producers also demonstrate a trade-off between stability and the ability of secondary producers to invade. Regardless of interclonal competition strength, increasing niche specificity promotes stability of the producer-only tumor such that cheaters are unable to invade. This result agrees with Gerlee and Anderson’s findings that selection for niche construction requires sufficient specificity, as specificity keeps cheaters from free-riding [30]. However, we find that niche construction specificity makes it less likely that secondary producers can invade a producer-only tumor. This stems from the fact that secondary producers do not produce the primary resource and therefore are also selected against due to specificity of the resource. On the other hand, the ability of global producers to invade this tumor does not depend on niche construction specificity since they also produce the local resource and benefit with the same efficiency as local producers. Instead, invasion is possible only under competition structure I and only if interclonal competition between global and secondary producers is weaker than intraclonal competition.

Our main result is identifying a trade-off between the arrival of mutations leading to metastasis and their invasion success. This trade-off may help explain the early metastasis hypothesis, which posits that metastasis is not necessarily a late event in the tumor history, but rather can occur while the tumor is still small. Many genetic and clinical studies support this view [36]. For example, evidence suggests that cells in metastases are genetically less progressed in terms of tumor progression than primary tumor cells at diagnosis [37, 38] and that metastases do not necessarily not come from large tumors [39]. Studies of breast cancer metastasis suggest that it can be an early event [40–44]. Similarly, it has been proposed that metastatic capacity stems from mutations acquired early in a tumor history [45], an idea supported by a recent analysis of tumor phylogenies that shows early genetic divergence of metastatic lineages [31]. These observations contradict the idea that cancer follows a linear progression in which late-stage primary tumors facilitate metastasis. However, the idea that metastasis is not a late event may be paradoxical because late primary tumors are larger and harbor more mutations that can lead to the genesis of metastatic lineages. Our results indicate that this paradox and the early metastasis phenomenon may potentially operate through a tension between mutant arrival and invasion caused by competition between local and pre-metastatic niche constructors late in a tumor history. Secondary or global producers must invade while the tumor is still small, and if they do the pre-metastatic niche will begin recruiting circulating tumor cells from an early time point. Otherwise, the pre-metastatic niche may remain unprepared, since larger, late primary tumors containing producers may be resistant to invasion by pre-metastatic niche constructors. Large tumors participate in metastasis as long as invasion occurred while the tumor was still small. Accordingly, empirical evidence suggests that construction of the pre-metastatic niche is the limiting factor for establishing secondary tumors, not dissemination of circulating tumor cells which is independent of tumor size [39] and occurs starting early on [46]. This is supported by the parallel progression model of cancer, in which frequently disseminated cancer cells rarely establish themselves [31, 47]. In short, our results showing a tension between the arrival of a mutation for pre-metastatic niche construction and its successful establishment support the idea that metastasis begins early and provide a potential explanation for a paradoxical aspect of nonlinear tumor progression. Our conclusion that the timing of metastasis is partially mediated through the timing of invasion by pre-metastatic niche constructors into the primary tumor can be validated if empirical analyses reveal that mutations causing pre-metastatic niche construction occur before the divergence of metastatic tumor lineages from the primary tumor.

One implication of our results is that if certain types of cancers may be resistant to metastasis even over long periods of time despite the accumulation of mutations. This happens if the tumor switches to a producer-only or coexistence state and with high niche-construction specificity and relatively high competition between different producer clones. Cancers that are not associated with metastasis are known since the work of Paget [23], who first proposed the “seed and soil” hypothesis. Our model suggests there could be an ecological, rather than genetic, explanation for the tendency of certain cancers to be less likely to metastasize. Resistance to invasion of metastatic subclones can be characteristic of particular cancers based on the typical cell types within the primary tumor cell population and the way they compete and use resources, rather than from a lack of necessary mutations.

Under competition structure III we find that in the coexistence quasi-equilibrium, reducing the growth rate of cheaters promotes the invasion of secondary producers. Chemotherapy is a method of targeting rapidly dividing cells and likely to disproportionately affect cheaters [48]. Thus, it is possible for chemotherapy to depress cheater growth rate enough such that *r*_00_ < *r*_01_, which would lead to equation 17 to be satisfied and secondary producers to invade, a necessary step towards metastasis. This result is consistent with accumulating evidence that chemotherapy may increase the potential for metastasis by increasing pro-tumorigenic growth factors in the blood and mobilizing bone marrow-derived progenitor cells to make the secondary tumor site more receptive to circulating tumor cells [49–51].

The assumption that intra-type competition is stronger than inter-type competition is central to our results. This assumption, shared with other models of clonal dynamics [52, 53], can be viewed as an expression of the fact that cellular neighbors tend to be of the same cell type and competition for resources occurs on a local spatial scale. A wealth of mathematical and experimental evidence shows that tumors contain spatial clustering of subclones with relatedness decreasing as distance increases between cells [30, 54–60]. Another biological mechanism for stronger intraclonal competition is that different producer and cheater clones might occupy different niches, due to their different metabolic needs and utilization of different cellular pathways.

Another assumption we made was that the benefit of niche construction is manifested by increasing carrying capacity of both producers and cheaters [30]. Cancer cells thrive at cellular densities considerably higher than that of normal host cells [61]. Increased carrying capacity due to niche construction can be achieved through many mechanisms; perhaps the most obvious is angiogenesis. Tumors often live in highly acidic microenvironments due to their increased glycolytic metabolism. Inducing vascularization delivers oxygen, clears metabolic waste products, provides nutrients, and provides growth factors. It has been established that tumors larger than 1-2 mm are supported by newly formed blood vessels through secretion of various angiogenic factors, including PDGF (platelet-derived growth factor), AngI, AngII, and VEGF [62, 63]. One model used tumor carrying capacity as a function of blood vessel density due to the importance of tumor-induced angiogenesis [62], and this is essentially carrying capacity as a function of niche construction. Another example is the release of autocrine factors by tumor cells, since this increases their ability to divide despite high cell density [30]. In vitro [64] and in vivo [65] studies have observed tumors with a subset of producers that contributed to overall population growth through the secretion of diffusable growth factors. This is evidence that a tumor can have producers and cheaters with an increasing carrying capacity.

In summary, we have created a mathematical model to study metastasis as an outcome of niche construction. Our results suggest that there exists a tension between mutant arrival and invasion. Tumors containing cheaters only are completely susceptible to invasion by all producer cell types while tumors containing producers can be resistant to invasion, dependent on competition strength and niche construction specificity. Our findings may help explain the early metastasis phenomenon and the observation that metastasis involves early mutations. We emphasize that successful metastasis requires a “double-hit” of the necessary genetic mutations and appropriate ecological conditions. Much research has focused on the genetic aspects of cancer initiation and progression, but this is insufficient if the context in which the genes exist and mutations arise is not considered [1, 4]. Paget’s “seed and soil” hypothesis is often invoked while studying metastasis [23]; our model shows that the analogy is more than evocative. Just as we need to consider the soil, sunlight, wind, and nearby flora and fauna to understand the germination of a seed, we also need to take the ecologist’s view to understand metastasis. Only then can we hope to stop the seed from spreading in the first place.

## Acknowledgements

We would like to acknowledge A. Brown and B. Morsky for helpful comments regarding the manuscript. J.Q. was supported by a summer stipend from the Roy and Diana Vagelos Scholars Program in the Molecular Life Sciences at the University of Pennsylvania.

## Supplementary Information

### SI-A Extended model

The model we present in the main text can be extended to include dynamics in the bloodstream and secondary tumor site. We now consider three distinct ecosystems, denoted by subscripts 1,2, and 3, respectively: the primary tumor site, the bloodstream, and a secondary site that receives metastatic cells to form a secondary tumor. Their cell populations are respectively *N*_1_, *N*_2_, and *N*_3_ and can include both producers and cheaters. Cells are given a subscript (*i, j*), where *i* ∈ {0,1} describes the ability to participate in niche construction (0 for cheaters and 1 for producers) and *j* ∈ {1,2,3} denotes which ecosystem the cell is in (1 for primary tumor *β* 2 for bloodstream, and 3 for secondary tumor) The number of each cell type is *n*_*i,j*_, and the growth rate is *r*_*i,j*_

**Figure SI-A.1:**
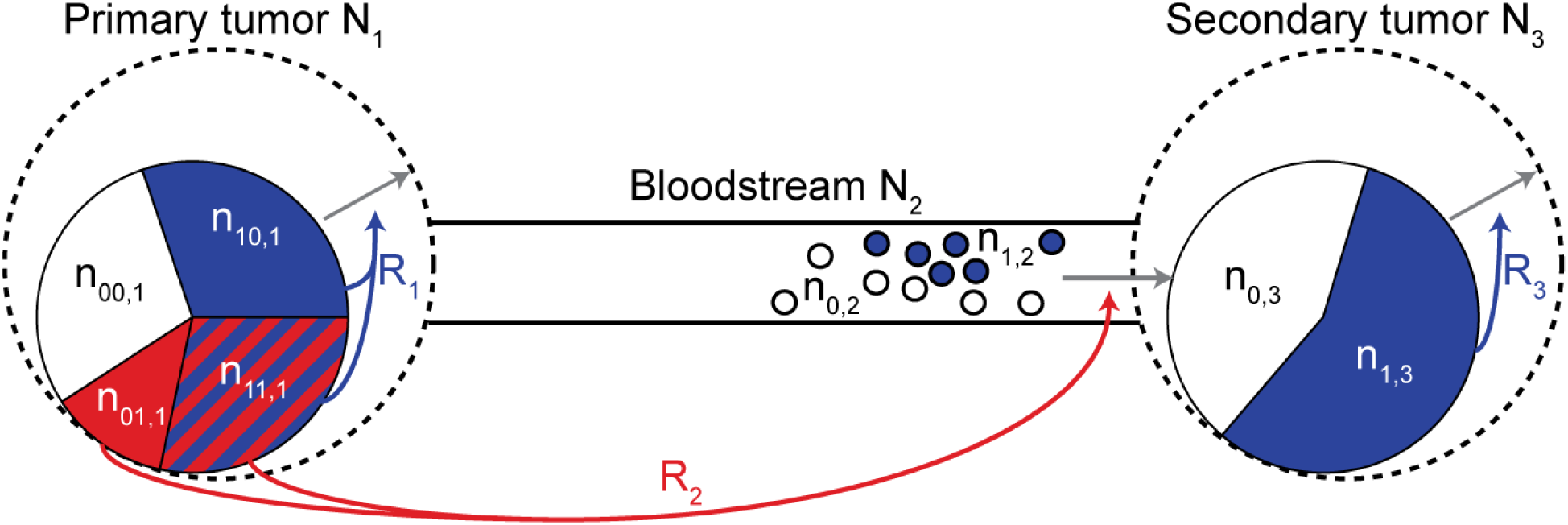
Schematic representation of the extended mathematical model. The model considers a primary tumor with four cell types, bloodstream with two cell types, and secondary tumor with two cell types. Cheaters are white and producers are blue. In the primary tumor, cells could additionally be secondary producers (red) or global producers (red and blue). Niche construction occurs in the tumor sites through production of resources *R*_1_ and *R*_3_, which benefit the tumors by increasing carrying capacity, represented as dotted lines. Construction of the pre-metastatic niche by primary tumor cells is represented by accumulation of resource *R*_2_, which facilitates settlement in the secondary tumor site.

*R*_1_ represents niche construction in the primary site. The extent to which the pre-metastatic niche has been constructed is measured by the amount of resource *R*_2_. For the primary tumor, *i* = *hk*, where *h* = 1 indicates local production and *k* = 1 indicates distant production. Local and global producers increase *R*_1_ with rate *g*_1_ while secondary and global producers increase *R*_2_ with rate *g*_2_. *R*_1_ and *R*_2_ suffer independent resource depletion rates of *l*_1_ and *l*_2_, respectively.

Primary tumor cells enter the bloodstream as a result of intravasation and die with rate *d* due to immune surveillance, anoikis, or physical stress. The intravasation function has the same form as discussed in the main text 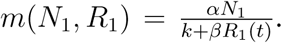 Though there is no niche construction while migrating through the bloodstream, cells retain their propensity for local niche construction local producers and global producers remain producers, while cheaters and secondary producers remain cheaters. We assume that once in the bloodstream, cells cease to engage in premetastatic niche construction, so we do not keep track of the distant producers and cheaters separately, reducing the number of cell types from four in the primary tumor to two in the bloodstream and secondary tumor: (00 1) and (01,1) cells become (0,2) cells in the bloodstream while (10,1) and (111) cells become (1,2) cells.

Cells can undergo extravasation and successfully settle the pre-metastatic niche as a function of *R*_2_. We assume that construction of the pre-metastatic niche is necessary for circulating tumor cells to settle down. In particular, we postulate a linear relationship between settlement rate and *R*_2_ with slope *δ*. Upon settling, (0,2) cells become (0,3) cells while (1,2) cells become (1,3) cells. There is local niche construction by (1,3) cells in this metastatic tumor which increases resource *R*_3_ with rate *g*_3_. *R*_3_ is a measure of local niche construction in the secondary tumor, which does not affect settlement but rather benefits cells that have already successfully metastasized. *R*_3_ also has independent resource depletion with rate *l*_3_.

**Table SI-A.1:** Governing equations of the extended model and the corresponding variables whose rates of change they describe, using competition structure I. Time dependence of *n* and *R* has been suppressed for notational simplicity. Dependence of *m* on *N*_1_ and *R*_1_ has also been suppressed.

|  |  |  |
| --- | --- | --- |
| Primary cheaters | $\frac{dn_{00,1}}{dt} = r_{00,1}n_{00,1} \left( 1 - \frac{n_{00,1} + \phi n_{01,1} + \theta n_{10,1} + \omega n_{11,1}}{k + \beta_0 R_1} \right) - mn_{00,1}$ | (SI-A.1) |
| Secondary producers | $\frac{dn_{01,1}}{dt} = r_{01,1}n_{01,1} \left( 1 - \frac{\phi n_{00,1} + n_{01,1} + \psi n_{10,1} + \nu n_{11,1}}{k + \beta_0 R_1} \right) - mn_{01,1}$ | (SI-A.2) |
| Primary producers | $\frac{dn_{10,1}}{dt} = r_{10,1}n_{10,1} \left( 1 - \frac{\theta n_{00,1} + \psi n_{01,1} + n_{10,1} + \mu n_{11,1}}{k + \beta_1 R_1} \right) - mn_{10,1}$ | (SI-A.3) |
| Global producers | $\frac{dn_{11,1}}{dt} = r_{11,1}n_{11,1} \left( 1 - \frac{\omega n_{00,1} + \nu n_{01,1} + \mu n_{10,1} + n_{11,1}}{k + \beta_1 R_1} \right) - mn_{11,1}$ | (SI-A.4) |
| Bloodstream cheaters | $\frac{dn_{0,2}}{dt} = \frac{\alpha N_1}{k + \beta_0 R_1} (n_{00,1} + n_{01,1}) - (d + \delta R_2)n_{0,2}$ | (SI-A.5) |
| Bloodstream producers | $\frac{dn_{1,2}}{dt} = \frac{\alpha N_1}{k + \beta_1 R_1} (n_{10,1} + n_{11,1}) - (d + \delta R_2)n_{1,2}$ | (SI-A.6) |
| Metastatic cheaters | $\frac{dn_{0,3}}{dt} = r_{0,3}n_{0,3} \left( 1 - \frac{n_{0,3} + \theta n_{1,3}}{k + \beta_0 R_3} \right) + \delta R_2 n_{0,2}$ | (SI-A.7) |
| Metastatic producers | $\frac{dn_{1,3}}{dt} = r_{1,3}n_{1,3} \left( 1 - \frac{\theta n_{0,3} + n_{1,3}}{k + \beta_1 R_3} \right) + \delta R_2 n_{1,2}$ | (SI-A.8) |
| Primary resource | $\frac{dR_1}{dt} = g_1 (n_{10,1} + n_{11,1}) - l_1 R_1$ | (SI-A.9) |
| Settlement resource | $\frac{dR_2}{dt} = g_2 (n_{01,1} + n_{11,1}) - l_2 R_2$ | (SI-A.10) |
| Metastasis resource | $\frac{dR_3}{dt} = g_3 n_{1,3} - l_3 R_3$ | (SI-A.11) |

### SI-B Alternative interpretation of cell types

Our model includes four cell types: cheaters, local producers, secondary producers, and global producers. Local and global producers contribute to primary niche construction, while secondary and global producers contribute to pre-metastatic niche construction. This interpretation of the cell populations can actually be generalized: as long as cells pay some cost to promote metastasis, whether it be via pre-metastatic niche construction or some other mechanism, the mathematical details and results of our model remain the same. This is because we focus on the prerequisites of metastasis within the primary tumor. For the extended model in SI section SI-A, the settlement dynamics would change based on the interpretation of cell types.

We present one potential alternative interpretation of the four cell types. Local producers pay a cost to participate in local niche construction benefiting all primary tumor cells, but have low metastatic potential. Cheaters benefit from local niche construction without paying the cost, and also possess low metastatic potential. The third cell type, analogous to the original secondary producer, benefits from local niche construction without paying the cost, but has high metastatic potential which comes at a cost. The fourth cell type, analogous to the original global producer, participates in local niche construction at a cost and also possesses high metastatic potential which comes at a cost. This interpretation focuses not on the construction of the pre-metastatic niche, but rather on metastatic potential of primary tumor cells, without changing any of the model’s mathematical details. In this framework, existence of cells with high metastatic potential is a prerequisite of metastasis. Metastatic potential can include various characteristics that promote the cell’s ability to successfully spawn a metastatic lesion, for example the ability to evade numerous cell death signals that are induced by loss of attachment to neighboring cells (anoikis) and the extracellular matrix (amorphosis) [1]. It is reasonable to assume high metastatic potential may incur a growth rate cost in the primary tumor. For example, the motile invasive phenotype, which fosters metastasis, may be characterized by a growth rate cost [2], which may stem from the fact that cells capable of moving cannot divide while moving [3, 4]. In short, our mathematical model is not sensitive to the specific interpretation of the cell types as long as there is a cost to promoting metastasis. In the main text, we focus on niche construction and the establishment of the pre-metastatic niche, but using other frameworks such as metastatic potential leads to the same results from the model.

### SI-C Competition structures

**Table SI-C.1:** Governing equations of the model for competition structure II.

|  |  |  |
| --- | --- | --- |
| Primary cheaters | $\frac{dn_{00}}{dt} = r_{00}n_{00}\left(1 - \frac{n_{00} + \phi n_{01} + \theta n_{10} + \psi n_{11}}{k + \beta_0 R}\right) - mn_{00}$ | (SI-C.1.1) |
| Secondary producers | $\frac{dn_{01}}{dt} = r_{01}n_{01}\left(1 - \frac{\phi n_{00} + n_{01} + n_{10} + n_{11}}{k + \beta_0 R}\right) - mn_{01}$ | (SI-C.1.2) |
| Primary producers | $\frac{dn_{10}}{dt} = r_{10}n_{10}\left(1 - \frac{\theta n_{00} + n_{01} + n_{10} + n_{11}}{k + \beta_1 R}\right) - mn_{10}$ | (SI-C.1.3) |
| Global producers | $\frac{dn_{11}}{dt} = r_{11}n_{11}\left(1 - \frac{\psi n_{00} + n_{01} + n_{10} + n_{11}}{k + \beta_1 R}\right) - mn_{11}$ | (SI-C.1.4) |
| Resource | $\frac{dR}{dt} = g(n_{10} + n_{11}) - lR$ | (SI-C.1.5) |

**Table SI-C.2:** Governing equations of the model for competition structure III.

Table SI-C.1: Governing equations of the model for competition structure II.
|  |  |  |
| --- | --- | --- |
| Primary cheaters | $\frac{dn_{00}}{dt} = r_{00}n_{00}\left(1 - \frac{n_{00} + n_{01} + \theta(n_{10} + n_{11})}{k + \beta_0 R}\right) - mn_{00}$ | (SI-C.2.1) |
| Secondary producers | $\frac{dn_{01}}{dt} = r_{01}n_{01}\left(1 - \frac{n_{00} + n_{01} + \theta(n_{10} + n_{11})}{k + \beta_0 R}\right) - mn_{01}$ | (SI-C.2.2) |
| Primary producers | $\frac{dn_{10}}{dt} = r_{10}n_{10}\left(1 - \frac{\phi(n_{00} + n_{01}) + n_{10} + n_{11}}{k + \beta_1 R}\right) - mn_{10}$ | (SI-C.2.3) |
| Global producers | $\frac{dn_{11}}{dt} = r_{11}n_{11}\left(1 - \frac{\phi(n_{00} + n_{01}) + n_{10} + n_{11}}{k + \beta_1 R}\right) - mn_{11}$ | (SI-C.2.4) |
| Resource | $\frac{dR}{dt} = g(n_{10} + n_{11}) - lR$ | (SI-C.2.5) |

### SI-D Supplementary simulations

**Figure SI-D.1:**
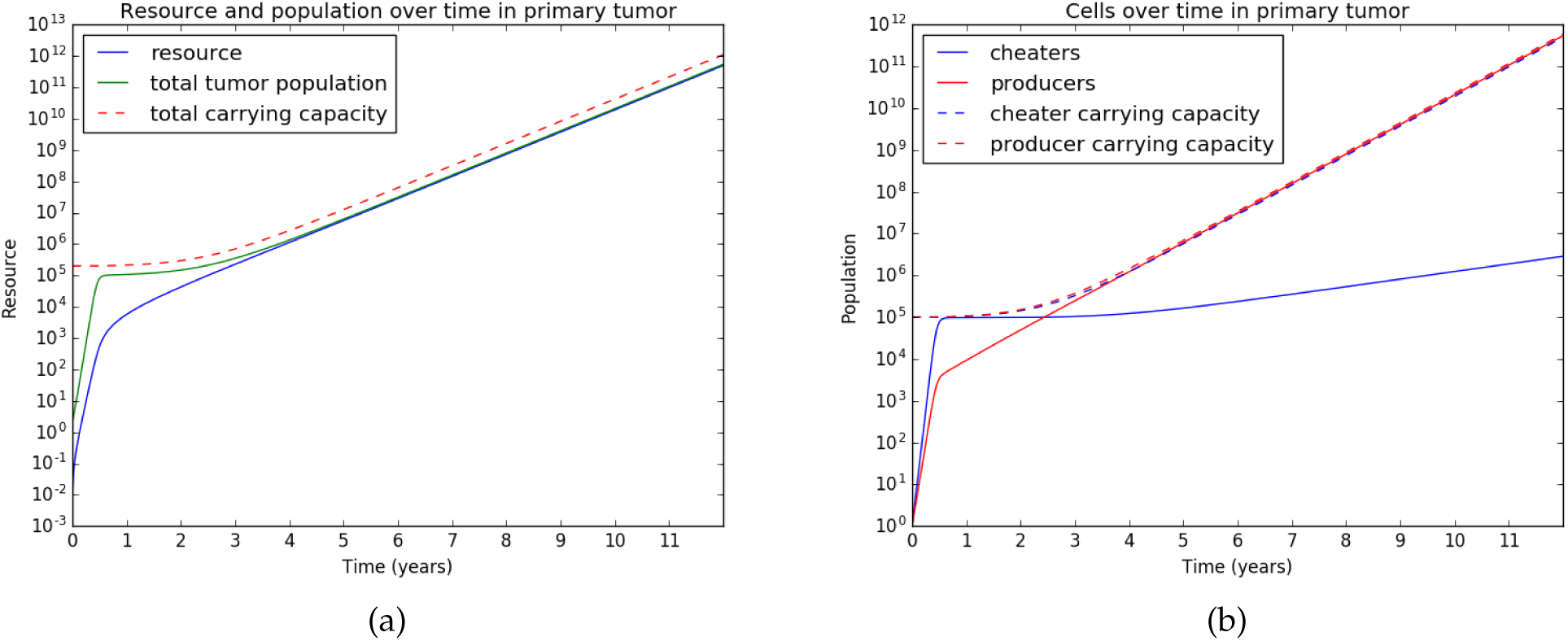
Simulation of a tumor with competition structure I starting with both cheaters and producers before the separation of time-scales. *r*_00_ = 0.07, *r*_10_ = 0.05, *r*_01_ = 0.045, *r*_11_ = 0.02, *k* = 10^5^,*β*_0_ = 1,*β*_1_ = 1.2, *θ* = 0.9, *g* = 0.004, *l* = 0.001, *α* = 10-^6^.

Figure SI-D.1 shows that prior to the separation of time-scales, the model (using competition structure I) contains a clinically realistic tumor size over time but fails to reach an equilibrium even after a decade. Different reasonable parameter combinations yield the same result. Cell populations in the model equilibrate more quickly than resource dynamics. The cell density always closely tracks the carrying capacity and the resource dynamics are slow. This allows us to make a separation of time-scales argument, which is biologically expected since niche construction processes (such as microenvironment vascularization) are generally slower than cell division.

### SI-E Estimation of parameters

In order to keep the model as realistic as possible, we attempted to choose parameters such that the simulated tumor reflects some behaviors observed clinically. According to one study, breast tumors grow on average 7.8 ± 3.9 years before detection, where the detection size is approximately 10^9^ cells [5] The value of *k*, the intrinsic carrying capacity, should reflect the fact that typical tumor population sizes can range from 106 to 10^11^ cells [6,7]. One estimate is that the size limit of cancer cell populations prior to the initiation of angiogenesis is 10^5^ [8], so this is a reasonable estimate for the intrinsic carrying capacity *k*. On the other hand, the size limit after angiogenesis is on the order of 10^12^ [8], which is generally viewed as the lethal tumor size at which patient death occurs [9]. Other methods estimate the maximum tumor size to be 12 cm [10] which corresponds to approximately 7.23 × 10^11^ cells [6], corroborating the 10^12^ estimate.

The intrinsic growth rates of clones can be estimated using data from the Norwegian Breast Cancer Screening Program [10]. The study used a logistic growth model and data from a large population to estimate doubling times of breast cancer tumors. In women aged 50-69 years, a 15 mm tumor doubled in diameter on average in 100 days while a 10 mm tumor doubled in diameter on average in 1.7 years. Using the conversion that one cubic centimeter tumor corresponds to 10^8^ cells [6], and assuming tumors are perfect spheres, these two doubling times can be converted to 0.0707 and 0.0113 day-^1^, taking into account size-dependent growth. Thus, a reasonable estimate for the highest growth rate r_00_ may be 0.07 day-^1^. This agrees with growth rates obtained from other studies using clinical data [9] and is similar to assumed parameters used in another mathematical model [11].

The rate of intravasation into the bloodstream has been estimated to be on the order of 10- to 10-^11^ day-^1^, but this seems to include the death rate of circulating cells [12]. It is often estimated that less than 1% of circulating tumor cells survive [13,14]. Another estimate of the integrated rate of leaving the primary site and successfully joining a secondary tumor in pancreatic cancer is 6 × 10 ^7^ per cell cycle [15]. We thus assume the parameter *β* describing intravasation is on the order of *α* = 10^−6^ day^−1^ or less, which is several orders of magnitudes smaller than all other parameters.

## References

[1] Basanta D, Anderson AR. Exploiting ecological principles to better understand cancer progression and treatment. Interface Focus 3 (2013), 20130020.

[2] Lloyd MC, Gatenby RA, Brown JS. “Ecology of the Metastatic Process”. In: Ecology and Evolution of Cancer. Elsevier, 2017, 153–165.

[3] Altrock PM, Liu LL, Michor F. The mathematics of cancer: integrating quantitative models. Nature Reviews Cancer 15 (2015), 730–745.

[4] Korolev KS, Xavier JB, Gore J. Turning ecology and evolution against cancer. Nature Reviews Cancer 14 (2014), 371–380.

[5] Laland KN, Odling-Smee FJ, Feldman MW. Evolutionary consequences of niche construction and their implications for ecology. Proceedings of the National Academy of Sciences 96 (1999), 10242–10247.

[6] Odling-Smee FJ, Laland KN, Feldman MW. Niche construction: the neglected process in evolution. Princeton University Press, 2003.

[7] Laland K, Matthews B, Feldman MW. An introduction to niche construction theory. Evolutionary Ecology 30 (2016), 191–202.

[8] Yang KR, Mooney SM, Zarif JC, Coffey DS, Taichman RS, Pienta KJ. Niche inheritance: a cooperative pathway to enhance cancer cell fitness through ecosystem engineering. Journal of Cellular Biochemistry 115 (2014), 1478–1485.

[9] Kareva I. Cancer ecology: Niche construction, keystone species, ecological succession, and ergodic theory. Biological Theory 10 (2015), 283–288.

[10] Bergman A, Gligorijevic B. Niche construction game cancer cells play. European Physical Journal Plus 130 (2015).

[11] Ibrahim-Hashim A, Gillies RJ, Brown JS, Gatenby RA. “Coevolution of Tumor Cells and Their Microenvironment:“Niche Construction in Cancer””. In: Ecology and Evolution of Cancer. Elsevier, 2017, 111–117.

[12] Hanahan D, Weinberg RA. Hallmarks of cancer: the next generation. Cell 144 (2011), 646–674.

[13] Wey J, Stoeltzing O, Ellis L. Vascular endothelial growth factor receptors: expression and function in solid tumors. Clinical Advances in Hematology & Oncology: H&O 2 (2004), 37–45.

[14] Catalano V, Turdo A, Di Franco S, Dieli F, Todaro M, Stassi G. “Tumor and its microenvironment: a synergistic interplay”. In: Seminars in Cancer Biology. Vol. 23. Elsevier. 2013, 522–532.

[15] Barar J, Omidi Y. Dysregulated pH in tumor microenvironment checkmates cancer therapy. BioImpacts: BI 3 (2013), 149.

[16] Pollak M. Insulin and insulin-like growth factor signalling in neoplasia. Nature Reviews Cancer 8 (2008), 915–928.

[17] Peinado H, Zhang H, Matei IR, Costa-Silva B, Hoshino A, Rodrigues G, Psaila B, Kaplan RN, Bromberg JF, Kang Y, et al. Pre-metastatic niches: organ-specific homes for metastases. Nature Reviews Cancer 17 (2017), 302–317.

[18] Peinado H, Lavotshkin S, Lyden D. “The secreted factors responsible for pre-metastatic niche formation: old sayings and new thoughts”. In: Seminars in Cancer Biology. Vol. 21. Elsevier. 2011, 139–146.

[19] Erler JT, Bennewith KL, Cox TR, Lang G, Bird D, Koong A, Le QT, Giaccia AJ. Hypoxia-induced lysyl oxidase is a critical mediator of bone marrow cell recruitment to form the premetastatic niche. Cancer Cell 15 (2009), 35–44.

[20] Peinado H, Alečković M, Lavotshkin S, Matei I, Costa-Silva B, Moreno-Bueno G, Hergueta-Redondo M, Williams C, GarcÍa-Santos G, Ghajar CM, et al. Melanoma exosomes educate bone marrow progenitor cells toward a pro-metastatic phenotype through MET. Nature medicine 18 (2012), 883–891.

[21] Psaila B, Lyden D. The metastatic niche: adapting the foreign soil. Nature Reviews Cancer 9 (2009), 285–293.

[22] Kaplan RN, Riba RD, Zacharoulis S, Bramley AH, Vincent L, Costa C, MacDonald DD, Jin DK, Shido K, Kerns SA, et al. VEGFR1-positive haematopoietic bone marrow progenitors initiate the pre-metastatic niche. Nature 438 (2005), 820–827.

[23] Paget S. The distribution of secondary growths in cancer of the breast. The Lancet 133 (1889), 571–573.

[24] Hiratsuka S, Watanabe A, Aburatani H, Maru Y. Tumour-mediated upregulation of chemoattractants and recruitment of myeloid cells predetermines lung metastasis. Nature Cell Biology 8 (2006), 1369–1375.

[25] Hiratsuka S, Watanabe A, Sakurai Y, Akashi-Takamura S, Ishibashi S, Miyake K, Shibuya M, Akira S, Aburatani H, Maru Y. The S100A8–serum amyloid A3–TLR4 paracrine cascade establishes a pre-metastatic phase. Nature Cell Biology 10 (2008), 1349–1355.

[26] Hood JL, San RS, Wickline SA. Exosomes released by melanoma cells prepare sentinel lymph nodes for tumor metastasis. Cancer research 71 (2011), 3792–3801.

[27] Jung T, Castellana D, Klingbeil P, Hernández IC, Vitacolonna M, Orlicky DJ, Roffler SR, Brodt P, Zöller M. CD44v6 dependence of premetastatic niche preparation by exosomes. Neoplasia 11 (2009), 1093IN13–1105IN17.

[28] Liu Y, Xiang X, Zhuang X, Zhang S, Liu C, Cheng Z, Michalek S, Grizzle W, Zhang HG. Contribution of MyD88 to the tumor exosome-mediated induction of myeloid derived suppressor cells. The American journal of pathology 176 (2010), 2490–2499.

[29] Grange C, Tapparo M, Collino F, Vitillo L, Damasco C, Deregibus MC, Tetta C, Bus-solati B, Camussi G. Microvesicles released from human renal cancer stem cells stimulate angiogenesis and formation of lung premetastatic niche. Cancer research 71 (2011), 5346–5356.

[30] Gerlee P, Anderson AR. The evolution of carrying capacity in constrained and expanding tumour cell populations. Physical Biology 12 (2015), 056001.

[31] Zhao ZM, Zhao B, Bai Y, Iamarino A, Gaffney SG, Schlessinger J, Lifton RP, Rimm DL, Townsend JP. Early and multiple origins of metastatic lineages within primary tumors. Proceedings of the National Academy of Sciences 113 (2016), 2140–2145.

[32] Alix-Panabières C, Riethdorf S, Pantel K. Circulating tumor cells and bone marrow micrometastasis. Clinical Cancer Research 14 (2008), 5013–5021.

[33] Pantel K, Brakenhoff RH. Dissecting the metastatic cascade. Nature Reviews Cancer 4 (2004), 448–456.

[34] Jackson AL, Loeb LA. The mutation rate and cancer. Genetics 148 (1998), 1483–1490.

[35] Cos S, Recio J, Sanchez-Barcelo E. Modulation of the length of the cell cycle time of MCF-7 human breast cancer cells by melatonin. Life Sciences 58 (1996), 811–816.

[36] Iskandar R. A Theoretical Model Of Breast Tumor Metastases In The Context Of Tumor Dormancy (2016).

[37] Klein CA, Hölzel D. Systemic cancer progression and tumor dormancy: mathematical models meet single cell genomics. Cell Cycle 5 (2006), 1788–1798.

[38] Schardt JA, Meyer M, Hartmann CH, Schubert F, Schmidt-Kittler O, Fuhrmann C, Polzer B, Petronio M, Eils R, Klein CA. Genomic analysis of single cytokeratin-positive cells from bone marrow reveals early mutational events in breast cancer. Cancer Cell 8 (2005), 227–239.

[39] Hüsemann Y, Geigl JB, Schubert F, Musiani P, Meyer M, Burghart E, Forni G, Eils R, Fehm T, Riethmüller G, et al. Systemic spread is an early step in breast cancer. Cancer Cell 13 (2008), 58–68.

[40] Braun S, Vogl FD, Naume B, Janni W, Osborne MP, Coombes RC, Schlimok G, Diel IJ, Gerber B, Gebauer G, et al. A pooled analysis of bone marrow micrometastasis in breast cancer. New England Journal of Medicine 353 (2005), 793–802.

[41] Schmidt-Kittler O, Ragg T, Daskalakis A, Granzow M, Ahr A, Blankenstein TJ, Kauf-mann M, Diebold J, Arnholdt H, Müller P, et al. From latent disseminated cells to overt metastasis: genetic analysis of systemic breast cancer progression. Proceedings of the National Academy of Sciences 100 (2003), 7737–7742.

[42] Bragado P, Sosa MS, Keely P, Condeelis J, Aguirre-Ghiso JA. “Microenvironments dictating tumor cell dormancy”. In: Minimal residual disease and circulating tumor cells in breast cancer. Springer, 2012, 25–39.

[43] Van’t Veer LJ, Dai H, Van De Vijver MJ, He YD, Hart AA, Mao M, Peterse HL, Van Der Kooy K, Marton MJ, Witteveen AT, et al. Gene expression profiling predicts clinical outcome of breast cancer. Nature 415 (2002), 530–536.

[44] Hosseini H, Obradović MM, Hoffmann M, Harper KL, Sosa MS, Werner-Klein M, Nanduri LK, Werno C, Ehrl C, Maneck M, et al. Early dissemination seeds metastasis in breast cancer. Nature 540 (2016), 552–558.

[45] Bernards R, Weinberg RA. Metastasis genes: a progression puzzle. Nature 418 (2002), 823–823.

[46] Eyles J, Puaux AL, Wang X, Toh B, Prakash C, Hong M, Tan TG, Zheng L, Ong LC, Jin Y, et al. Tumor cells disseminate early, but immunosurveillance limits metastatic outgrowth, in a mouse model of melanoma. The Journal of Clinical Investigation 120 (2010), 2030.

[47] Klein CA. Parallel progression of primary tumours and metastases. Nature Reviews Cancer 9 (2009), 302–312.

[48] Archetti M. Evolutionary game theory of growth factor production: implications for tumour heterogeneity and resistance to therapies. British Journal of Cancer 109 (2013), 1056–1062.

[49] Daenen LG, Roodhart JM, Amersfoort M van, Dehnad M, Roessingh W, Ulfman LH, Derksen PW, Voest EE. Chemotherapy enhances metastasis formation via VEGFR-1–expressing endothelial cells. Cancer Research 71 (2011), 6976–6985.

[50] Roodhart JM, Langenberg MH, Vermaat JS, Lolkema MP, Baars A, Giles RH, Witteveen EO, Voest EE. Late release of circulating endothelial cells and endothelial progenitor cells after chemotherapy predicts response and survival in cancer patients. Neoplasia 12 (2010), 87–94.

[51] Shaked Y, Henke E, Roodhart JM, Mancuso P, Langenberg MH, Colleoni M, Daenen LG, Man S, Xu P, Emmenegger U, et al. Rapid chemotherapy-induced acute endothelial progenitor cell mobilization: implications for antiangiogenic drugs as chemosensitizing agents. Cancer Cell 14 (2008), 263–273.

[52] Krepkin K, Costa J. Defining the role of cooperation in early tumor progression. Journal of Theoretical Biology 285 (2011), 36–45.

[53] Krepkin K. The Role of Cooperation in Pre-tumor Progression: A Cellular Population Dynamics Model (2010).

[54] González-Garcìa I, Solé RV, Costa J. Metapopulation dynamics and spatial heterogeneity in cancer. Proceedings of the National Academy of Sciences 99 (2002), 13085–13089.

[55] Nakamura T, Kuwai T, Kitadai Y, Sasaki T, Fan D, Coombes KR, Kim SJ, Fidler IJ. Zonal heterogeneity for gene expression in human pancreatic carcinoma. Cancer Research 67 (2007), 7597–7604.

[56] Greaves M, Maley CC. Clonal evolution in cancer. Nature 481 (2012), 306–313.

[57] Maley CC, Galipeau PC, Finley JC, Wongsurawat VJ, Li X, Sanchez CA, Paulson TG, Blount PL, Risques RA, Rabinovitch PS, et al. Genetic clonal diversity predicts progression to esophageal adenocarcinoma. Nature Genetics 38 (2006), 468–473.

[58] Yachida S, Jones S, Bozic I, Antal T, Leary R, Fu B, Kamiyama M, Hruban RH, Eshleman JR, Nowak MA, et al. Distant metastasis occurs late during the genetic evolution of pancreatic cancer. Nature 467 (2010), 1114–1117.

[59] Clark J, Attard G, Jhavar S, Flohr P, Reid A, De-Bono J, Eeles R, Scardino P, Cuzick J, Fisher G, et al. Complex patterns of ETS gene alteration arise during cancer development in the human prostate. Oncogene 27 (2008), 1993–2003.

[60] Navin N, Krasnitz A, Rodgers L, Cook K, Meth J, Kendall J, Riggs M, Eberling Y, Troge J, Grubor V, et al. Inferring tumor progression from genomic heterogeneity. Genome Research 20 (2010), 68–80.

[61] Neri A, Welch D, Kawaguchi T, Nicolson GL. Development and Biologic Properties of Malignant Cell Sublines and Clones of a Spontaneously Metastasizing Rat Mammary Adenocarcinoma 2 3. Journal of the National Cancer Institute 68 (1982), 507–517.

[62] Bodnar M, Foryś U. Angiogenesis model with carrying capacity depending on vessel density. Journal of Biological Systems 17 (2009), 1–25.

[63] Folkman J. Angiogenesis in cancer, vascular, rheumatoid and other disease. Nature Medicine 1 (1995), 27–30.

[64] Archetti M, Ferraro DA, Christofori G. Heterogeneity for IGF-II production maintained by public goods dynamics in neuroendocrine pancreatic cancer. Proceedings of the National Academy of Sciences 112 (2015), 1833–1838.

[65] Marusyk A, Tabassum DP, Altrock PM, Almendro V, Michor F, Polyak K. Non-cell autonomous tumor-growth driving supports sub-clonal heterogeneity. Nature 514 (2014), 54.

## References cited in the Supplementary Information

[1] Mehlen P, Puisieux A. Metastasis: a question of life or death. Nature Reviews Cancer 6 (2006), 449–458.

[2] Basanta D, Hatzikirou H, Deutsch A. Studying the emergence of invasiveness in tumours using game theory. The European Physical Journal B-Condensed Matter and Complex Systems 63 (2008), 393–397.

[3] Giese A, Loo MA, Tran N, Haskett D, Coons SW, Berens ME. Dichotomy of astrocytoma migration and proliferation. International journal of cancer 67 (1996), 275–282

[4] Giese A, Bjerkvig R, Berens M, Westphal M. Cost of migration: invasion of malignant gliomas and implications for treatment. Journal of clinical oncology 21 (2003), 1624–1636.

[5] Speer JF, Petrosky VE, Retsky MW, Wardwell RH. A stochastic numerical model of breast cancer growth that simulates clinical data. Cancer Research 44 (1984), 4124–4130.

[6] Del Monte U. Does the cell number 109 still really fit one gram of tumor tissue? Cell Cycle 8 (2009), 505–506.

[7] Lipinski KA, Barber LJ, Davies MN, Ashenden M, Sottoriva A, Gerlinger M. Cancer evolution and the limits of predictability in precision cancer medicine. Trends in Cancer 2 (2016), 49–63.

[8] Clare SE, Nakhlis F, Panetta JC. Molecular biology of breast metastasis The use of mathematical models to determine relapse and to predict response to chemotherapy in breast cancer. Breast Cancer Research 2 (2000), 430.

[9] Spratt JA, Von Fournier D, Spratt JS, Weber EE. Decelerating growth and human breast cancer. Cancer 71 (1993), 2013–2019.

[10] Weedon-Fekjær H, Lindqvist BH, Vatten LJ, Aalen OO, Tretli S. Breast cancer tumor growth estimated through mammography screening data. Breast Cancer Research 10 (2008), R41.

[11] Leder K, Holland EC, Michor F. The therapeutic implications of plasticity of the cancer stem cell phenotype. PloS One 5 (2010), e14366.

[12] Iskandar R. A Theoretical Model Of Breast Tumor Metastases In The Context Of Tumor Dormancy (2016).

[13] Yang KR, Mooney SM, Zarif JC, Coffey DS, Taichman RS, Pienta KJ. Niche inheritance: a cooperative pathway to enhance cancer cell fitness through ecosystem engineering. Journal of Cellular Biochemistry 115 (2014), 1478–1485.

[14] Fidler I. Selection of successive tumour lines for metastasis. Nature 242 (1973), 148–149.

[15] Haeno H, Gonen M, Davis MB, Herman JM, Iacobuzio-Donahue CA, Michor F. Computational modeling of pancreatic cancer reveals kinetics of metastasis suggesting optimum treatment strategies. Cell 148 (2012), 362–375.

